# Predictive Control of Musculotendon Loads Across Fast and Slow-twitch Muscles in a Simulated System with Parallel Actuation

**DOI:** 10.1101/2024.05.14.594110

**Authors:** Mahdi Nabipour, Gregory S. Sawicki, Massimo Sartori

## Abstract

Research in lower limb wearable robotic control have largely focused on reducing the metabolic cost of walking or compensating for a portion of the biological joint torque e.g., by applying support proportional to estimated biological joint torques. However, due to different musculotendon unit (MTU) contractile speed properties, less attention has been given to the development of wearable robotic controllers that can steer MTU dynamics directly. Therefore, closed-loop control of MTU dynamics needs to be robust across fiber phenotypes, i.e. ranging from slow type I to fast type IIx in humans. The ability to closed-loop control the in-vivo dynamics of MTUs, could lead to a new class of wearable robots that can provide precise support to targeted MTUs, for preventing onset of injury or to provide precision rehabilitation to selected damaged tissues. In this paper, we introduce a novel closed-loop control framework that utilizes Nonlinear Model Predictive Control (NMPC) to keep the peak Achilles tendon force within predetermined boundaries during diverse range of cyclic force production simulations in the human ankle plantarflexors. This control framework employs a computationally efficient model comprising a modified Hill-type MTU contraction dynamics component and a model of the ankle joint with parallel actuation. Results indicate that the closed-form muscle-actuation model’s computational time is in the order of microseconds and is robust to different muscle contraction velocity properties. Furthermore, the controller achieves tendon force control within a timeframe below 14 *ms*, aligning with the physiological electromechanical delay of the MTU and facilitating its potential for future real-world applications.

## 1. Introduction

Recent effort in lower limb wearable exoskeleton and exosuit control has largely focused on two main objectives: reducing metabolic energy expenditure during locomotion Zhang et al. (2017); Witte et al. (2020); Slade et al. (2022) and compensating for a portion of biological joint torque via e.g., neuromus-culoskeletal (NMS) models, or regression techniques Durandau et al. (2022). However, the development of wearable robots capable of gaining direct closed-loop control over the mechanical loads acting on individual muscle-tendon units (MTUs) has not been addressed yet. The ability of gaining direct control on the loads acting on individual skeletal muscles or on series elastic tendons would enable wearable robots to provide bio-protective support e.g., by computing exoskeleton’s optimal support torques that prevent the risk of musculoskeletal injuries or tissue maladaptation linked with repetitive intense loads leading to tissue catabolic function. Similarly, this new class of wearable robots, could deliver precise mechanical stress and strain to injured tissues to facilitate stress- and strain-induced tissue regeneration following injury, opening to precision robot-aided rehabilitation.

Gaining direct control on individual MTU dynamics is however complex. MTUs display a large repertoire of dynamic behaviors, which is in part determined by the constituent muscle fiber phenotype Wisdom et al. (2015). Skeletal muscle fibers’s sarcomeres can express slow (i.e., type I) to fast (i.e., type II-a and II-x) myosin heavy chain isoforms, impacting the the ability of the whole muscle to generate force per unit time.

The closed-loop control of muscles underlying different contractile speed characteristics requires a robust framework, never achieved so far. There currently are no wearable exoskeletons and exosuits capable of predicting how external loads impact the load on biological tendons. Moreover, there is currently no robot capable of controlling the biological MTU load within predefined limits, regardless of muscle fibers phenotype, locomotion speed, frequency and duty cycle.

Recently, one study implemented nine pre-established plantarflexion assistance profiles to control Achilles tendon force within an ankle exosuit Schmitz et al. (2022). Additionally, Nuckols et al. devised an ankle torque profile based on ultrasound measurements of soleus contraction velocity during walking, aiming to minimize metabolic cost during costly concentric contractions Nuckols et al. (2021). Notably, the required processing time for this study was considerably shorter that the fastest human-in-the-loop (HIL) optimization used for minimising metabolic expenditure of walking Slade et al. (2022).

However, existing control schemes provide assistive torques to biological joints with no knowledge on the effect on joint’s constituent MTU force. Here we present an initial step towards addressing this gap by creating a predictive model-based control framework for closed-loop control of peak MTU load during cyclic motions such as hopping. The ideal controller for this application should obtain the assistive profile within milliseconds and should be able to predict the impact of the assistive load on the MTU mechanics simultaneously. To achieve this, we employ Nonlinear Model Predictive Control (NMPC) within the control framework with the aim of controlling the peak Achilles tendon force using a simulated exoskeleton parallel to the ankle plantarflexors during cyclic motions, such as hopping and walking.

For the model-based predictive control framework we obtain a combined MTU-exoskeleton system in ordinary differential equation (ODE) form. Hill-type models are commonly utilized to model MTU dynamics, primarily for their efficiency and simplicity in parameterization when compared to alternatives such as Huxley’s models of muscle contraction Huxley & Niedergerke (1954); Huxley & Hanson (1954). We utilize the conventional Hill-type model with an additional damping element parallel to the contractile element. In the existing literature, implicit De Groote et al. (2016) and numerical solutions Millard et al. (2013) for Hill-type models can be found.

In a prior preliminary study, we employed univariate linear regression to develop a closed-form solution for the damped Hill-type muscle model Nabipour et al. (2023); Nabipour & Sartori (2023). Nevertheless, the resulting model did not exhibit generalizability across various locomotion speeds and different muscle types. Consequently, we propose a computationally-efficient simplified closed-form solution for the damped Hill-type model to be integrated into the predictive control framework.

The contractile element of a Hill-type muscle encompasses a force-velocity relationship, which we utilize to model slow to fast-twitch muscle fiber phenotypes. A sensitivity analysis will be undertaken to assess the controller’s responsiveness to closed-loop regulation of peak Achilles tendon force during simulated human hopping, with varying levels of muscle excitation amplitudes. This analysis will also consider different control horizons for the NMPC and levels of support within the control framework. Additionally, we will investigate the controller’s robustness in controlling peak tendon force across muscles with differing twitch properties.

## 2. Methods

Below, the process of deriving the closed-form set of ODEs for the combined MTU-exoskeleton system, with a focus on hopping motion as a representative cyclic locomotion-like movement, is explained in detail. Subsequently, an analysis of the control structure and the methodology employed for assessing the robustness and sensitivity of the control framework will be presented.

### 2.1. Modeling combined MTU-exoskeleton system

The simulation of hopping motion involves simplifying the combined MTU-exoskeleton system into two main components: a lumped plantarflexor muscle (representing the whole triceps surae) and a parallel exoskeleton actuator capable of providing assistance parallel to the plantarflexors (Figure 1). This simplification assumes the presence of a single lumped MTU instead of individual plantarflexing MTUs Robertson et al. (2014); Robertson & Sawicki (2014).

**Figure 1.**
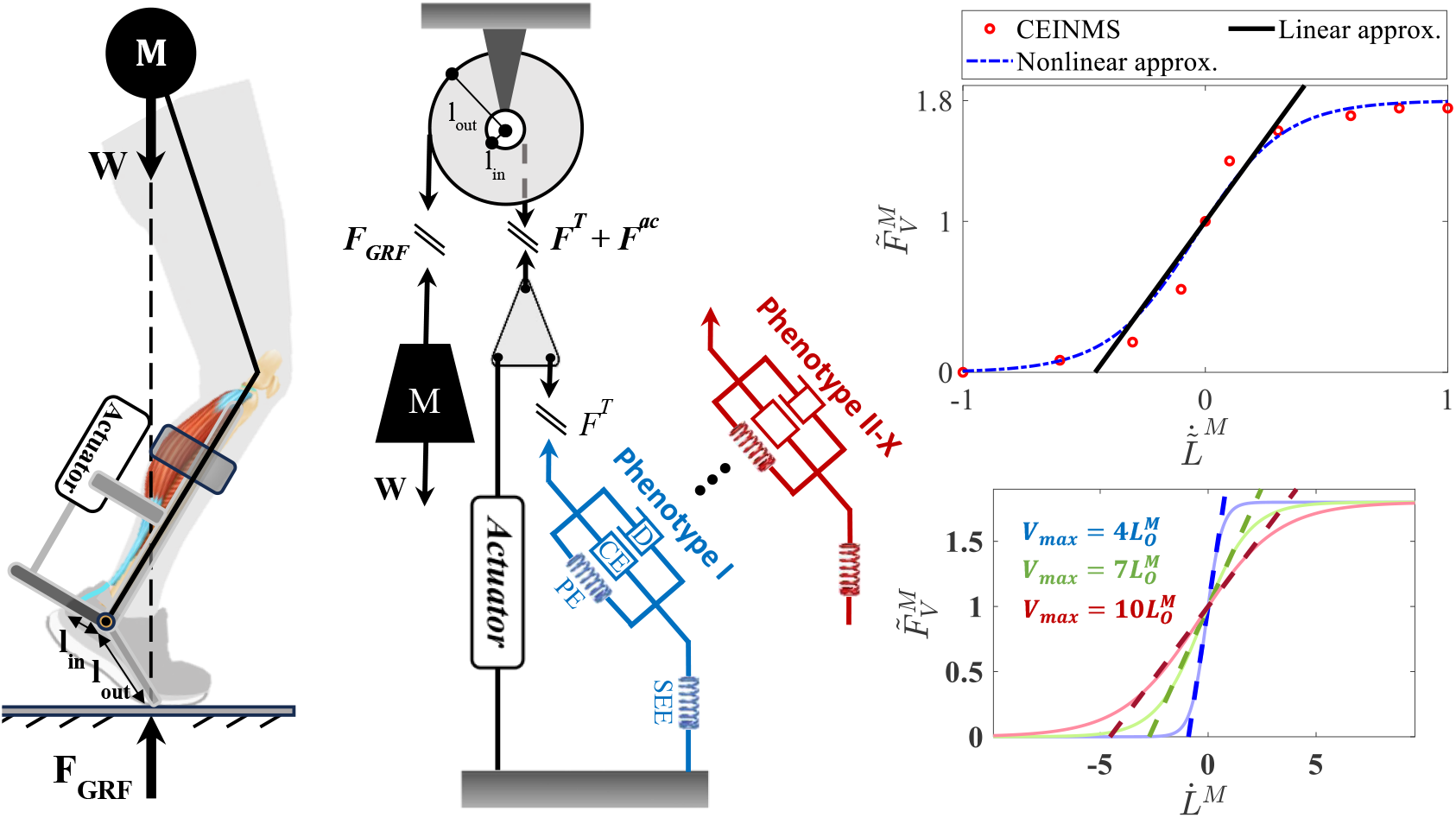
From left to right, the human-exoskeleton system representing the hopping model, the simplified pulley-mass model of hopping, and the velocity dependent F-V relation of the MTU contractile element.

#### 2.1.1. Modeling lumped triceps surae

We utilize a Hill-type model incorporating a pennation angle to simulate MTU dynamics Caillet et al. (2022); Thelen (2003). This model conventionally comprises the contractile element of the muscle arranged in both series and parallel configurations with elastic elements. We propose a modified version of this model where an additional damping element is added parallel to the muscle’s contractile and parallel elastic element, see Figure 1. As a result, the tendon force is derived as:

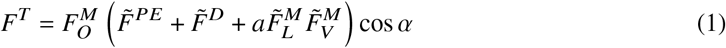

where 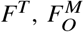 and *a*, and *α* represent the tendon force, maximum isometric force, muscle activation and pennation angle, respectively. Also, 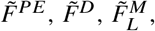, and 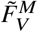 denote the normalized force of the parallel passive element, parallel damping, and contractile element’s active force-length (F-L) and force-velocity (F-V) relation. 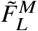 and 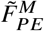 are derived from Anderson (2007). Damping’s governing equation 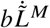 where b is the damping coefficient and 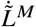 represents the normalized muscle fiber velocity.

To enable future predictions with a closed-form equation, it’s often advantageous to model the system ODEs:

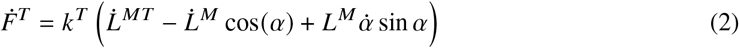

where, *k*^*T*^ and *L*^*M*^ are the tendon stiffness and muscle fiber length, respectively. The stiffness can either be derived from De Groote et al. (2016) or by ploynomial fit to stiffness or tendon force-strain data points used in our previously developed Calibrated EMG-Informed Neuromusculoskeletal Modelling (CEINMS) tool-box Pizzolato et al. (2015). 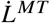 in equation 2 is directly derived from the kinematics of the movement. The 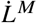 is often obtained via regression in the cases where the damping is not incorporated in the Hill-type model. When damping is considered, no closed-form solutions were derived and researchers often use numerical solutions Millard et al. (2013). The normalized muscle fiber velocity does not cross the range of 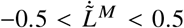 for walking and running with various speeds Arnold et al. (2013). Therefore, we simplify the F-V relation assuming muscle fiber contraction velocity remains within the linear range, as depicted in Figure 1:

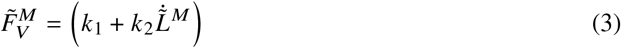

where, the optimal values for *k*_1_ and *k*_2_ that fits the F-V curve within CEINMS toolbox are 2.2 and 1, respectively. By combining equations 1 and 2, we can obtain:

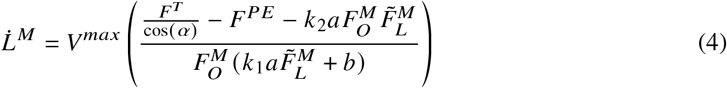

where *V* ^*max*^ represents the maximum fiber velocity which can take the range between 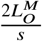 and 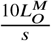 depending on whether the muscle’s constituent fiber phenotype is slow or fast-twitch, respectively. By replacing 3 in 2, the MTU contraction dynamics in ODE form will be obtained.

#### 2.1.2. Simplified model for hopping

The MTU contraction dynamics model should be combined with a motion-related equation to obtain the combined MTU-exoskeleton system. We use the same model used by Robertson et al. (2014); Robertson & Sawicki (2014) and add a parallel damping to the MTU contractile element. Additionally, we incorporate a parallel actuator into the system to simulate the exoskeleton.

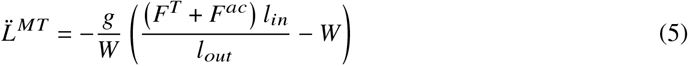

where the ratio 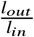 represent the mechanical advantage of the pulley-mass system shown in Figure 1. The combined MTU-exoskeleton model (equations 2 and 5) are used as the main model (the control plant) and within the control framework.

### 2.2 Tendon force control using NMPC

The objective of the NMPC algorithm is to minimize the following cost function while ensuring that the tendon force remains below a predetermined threshold:

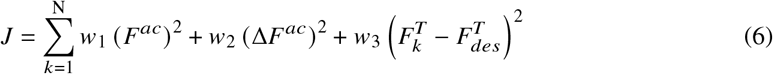

here, *N* represents the control horizon, while the weights *w*_*i*_ determine the influence of tendon force, actuator force (*F* ^*ac*^), and its increment (Δ*F*^*ac*^) on the overall cost function. We use a decreasing horizon and the smallest permissible horizon in our simulation is considered to be equal to 50 *ms* Nabipour et al. (2023). In all our simulations, we operate under the assumption that we have foreknowledge of the future MTU activation. This assumption remains valid particularly in cyclic tasks like hopping, where muscle synergies enable us to infer the activation throughout the entire cycle.

### 2.3 Control framework validation procedures

#### 2.3.1. Different horizons and levels of support

The initial prediction control horizon determines the duration over which the control framework predicts future MTU force profiles to decide whether to activate the NMPC. Additionally, as previously discussed, predicted future MTU force also serves as the starting point for the decreasing control horizon of the NMPC algorithm. Therefore, an excessively large initial control horizon can lead to significant time consumption, while overly small values may impair the controller’s ability to maintain the MTU load below a predetermined threshold. To analyze the sensitivity of the NMPC to this control parameter, we will simulate several initial horizons ranging from 50 to 125 *ms*, with an increment of 25 *ms*.

Having a controller robust to changes in levels of support, ensures the adaptability of the exoskeleton to different environments and enhances user’s comfort and safety. We will simulate variations in support levels by maintaining a fixed initial horizon and adjusting the threshold of the NMPC algorithm.

#### 2.3.2 Robustness to various muscle phenotypes

The investigation into the robustness of the control framework to changes in muscle fiber phenotypes is conducted in two ways: a) when the controller is informed of the phenotype change, and b) when the controller is unaware of the change. In all scenarios, a constant tendon force threshold of 2500 *N* is maintained. When the controller is aware of the phenotype change the *V* ^*max*^ in the hopping model and the controller have the same value. It should be noted that the tendon force profile changes due to change in the phenotype of the muscle fiber. Therefore, in figure 5 for every assisted case (solid lines) a non-assisted tendon force profile (dash-dotted lines) is presented.

Therefore, we simulate various fast and slow-twitch muscle phenotypes by tuning the parameter *p*_*i*_ in the equation 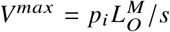 within the MTU model. In our simulations, we adjust this parameter across values of 4 (indicating slow-twitch muscle fiber), 7, and 10 (indicating fast-twitch muscle fiber). The controller’s ability to handle various phenotypes will be examined by simulating its behavior during hopping at a frequency of 2.5 *Hz*, both when the controller is aware and unaware of the change in muscle fiber phenotype. In the cases that the controller is aware of the change, the *V* ^*max*^ in the hopping model (control plant) and the framework (the future predictor and the NMPC inner model) are the same.

## 3. Results

### 3.1. Modeling

When simulating MTU dynamics using the conventional non-damped Hill-type muscle model, the tendon force solution becomes unstable whenever muscle activation approaches zero. Incorporating damping parallel to the contractile element and linearizing the F-V relation to derive a closed-form analytical solution not only resolves the instability issue but also reduces the Root Mean Square Error (RMSE) to 7.9% for simulating plantar and dorsiflexion movements on a dynamometer, see Figure 2.

**Figure 2.**
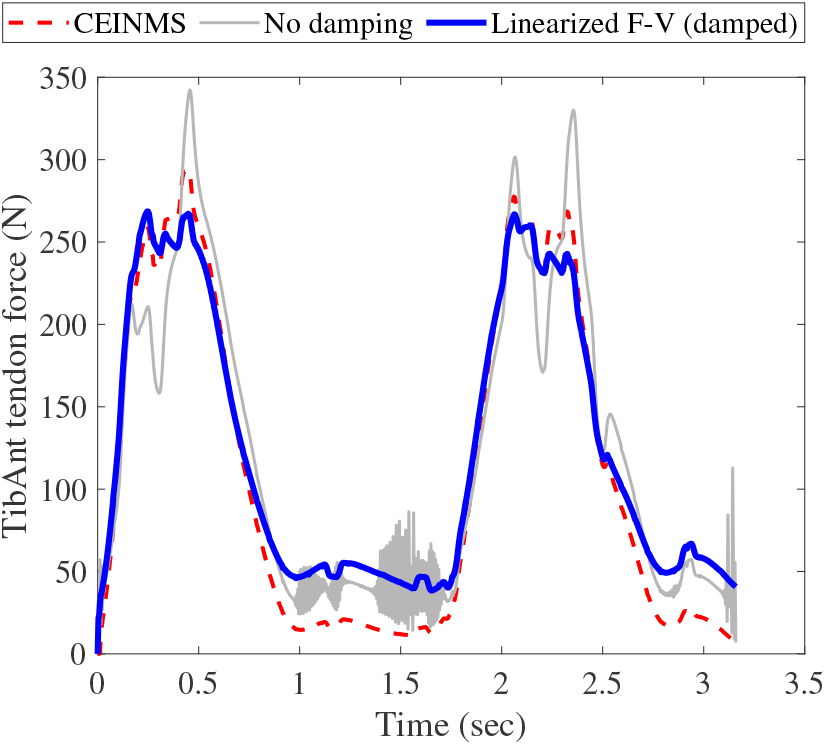
The effect of having parallel to the MTU contractile element and model’s accuracy when compared to the data collected on the dynamometer.

By changing the muscle fiber phenotype from type I to type II-X (*p*_*i*_ = 4, 7, 10) for Tibialis Anterior muscle during plantar and dorsiflexion movements on a dynamometer, we can see that the RMSE when changing the maximum velocity of the muscle from 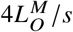 to 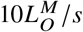 changes from 10.4% to 7.9%.

### 3.2 Predictive control

#### 3.2.1. Different horizons and levels of support

Since in our framework the horizon is varying, the initial horizon is of great significance. In Figure 3 the NMPC perfomance for different initial horizons starting from 50 *ms* up to 125 *ms* with increments of 25 *ms* is depicted.

**Figure 3.**
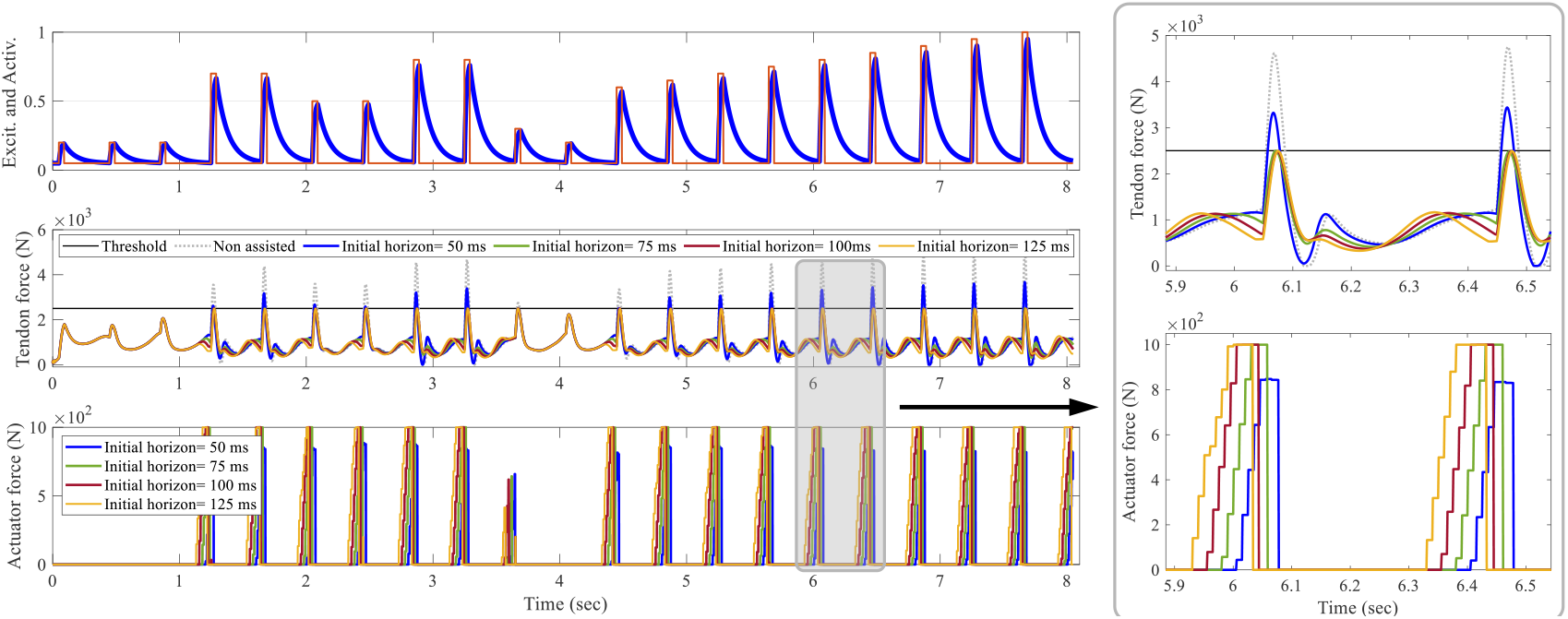
NMPC performance for different initial control horizons.

The maximum tendon force during simulation of hopping at a frequency of 2.5 *Hz* without assistance occurs when the muscle excitation amplitude is set to 1. In this scenario, the maximum tendon force reaches 5093 *N*. However, when assistance is introduced with a 50 *ms* initial horizon, the maximum tendon force reduces to 3671 *N*. The RMSE of the maximum tendon forces achieved in every hop is calculated to be 28.3%. The median and standard deviation (STD) of the maximum tendon forces are 3181.5 *N* and 405 *N*, respectively. For the initial horizon of 75 *ms*, the controller always keeps the maximum tendon force under the 2500 *N* threshold except in the case of the maximum excitation amplitude. In the cases of 100 *ms* and 125 *ms*, the controller always kept the maximum tendon force under the threshold. The maximum, median, and STD values for 100 *ms* (125 *ms*) are 2494 *N* (2494 *N*), 2474 *N* (2485 *N*), and 12.3% (11%), respectively.

The controller’s robustness to support different levels of assistance is also investigated, as shown in Figure 4. To demonstrate various levels of support, the threshold of the NMPC algorithm is adjusted every few hop, with an initial prediction horizon of 100*ms*. As depicted in Figure 4, the controller dynamically adjusts the control output to ensure that the peak tendon force remains below the threshold, achieving an RMSE of 35.7*N*.

**Figure 4.**
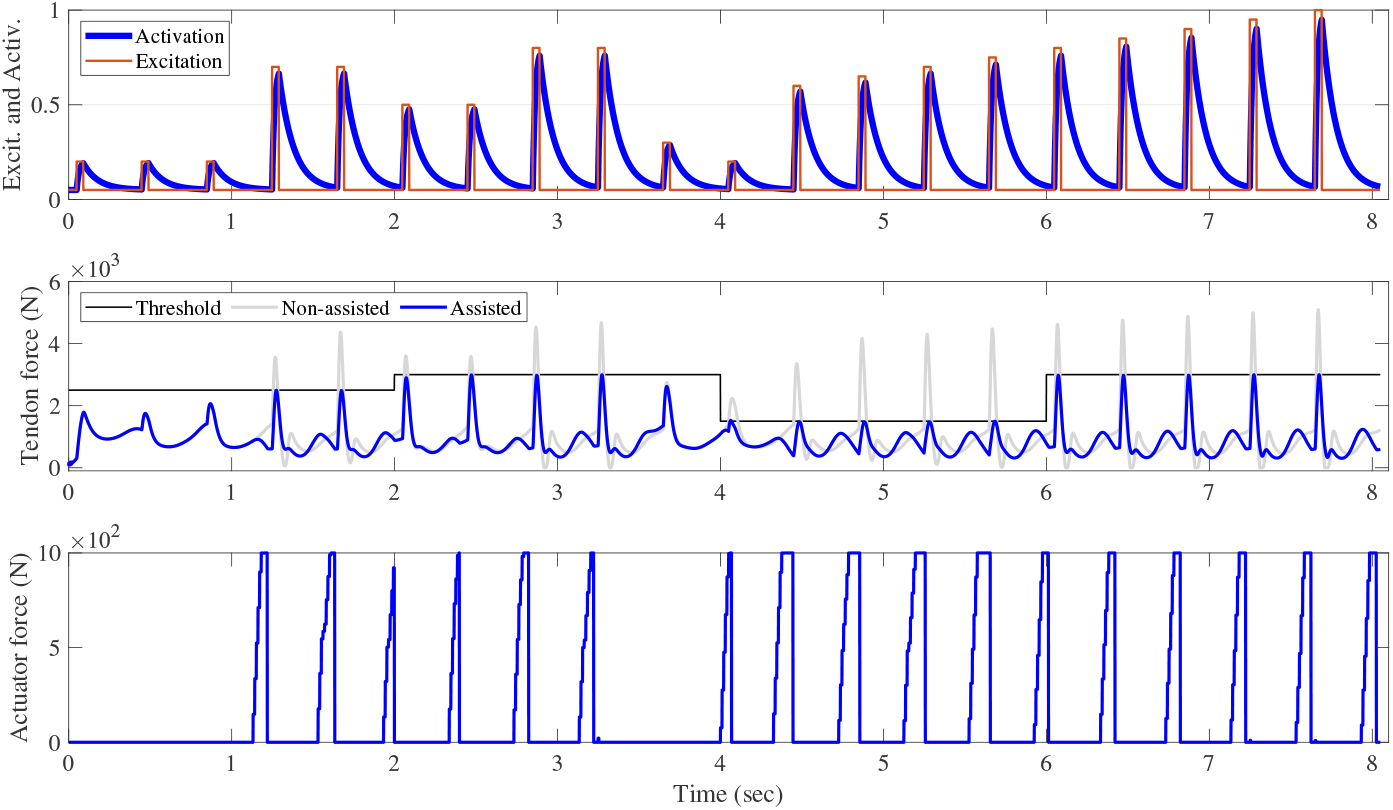
Control framework’s performance for different levels of support.

**Figure 5.**
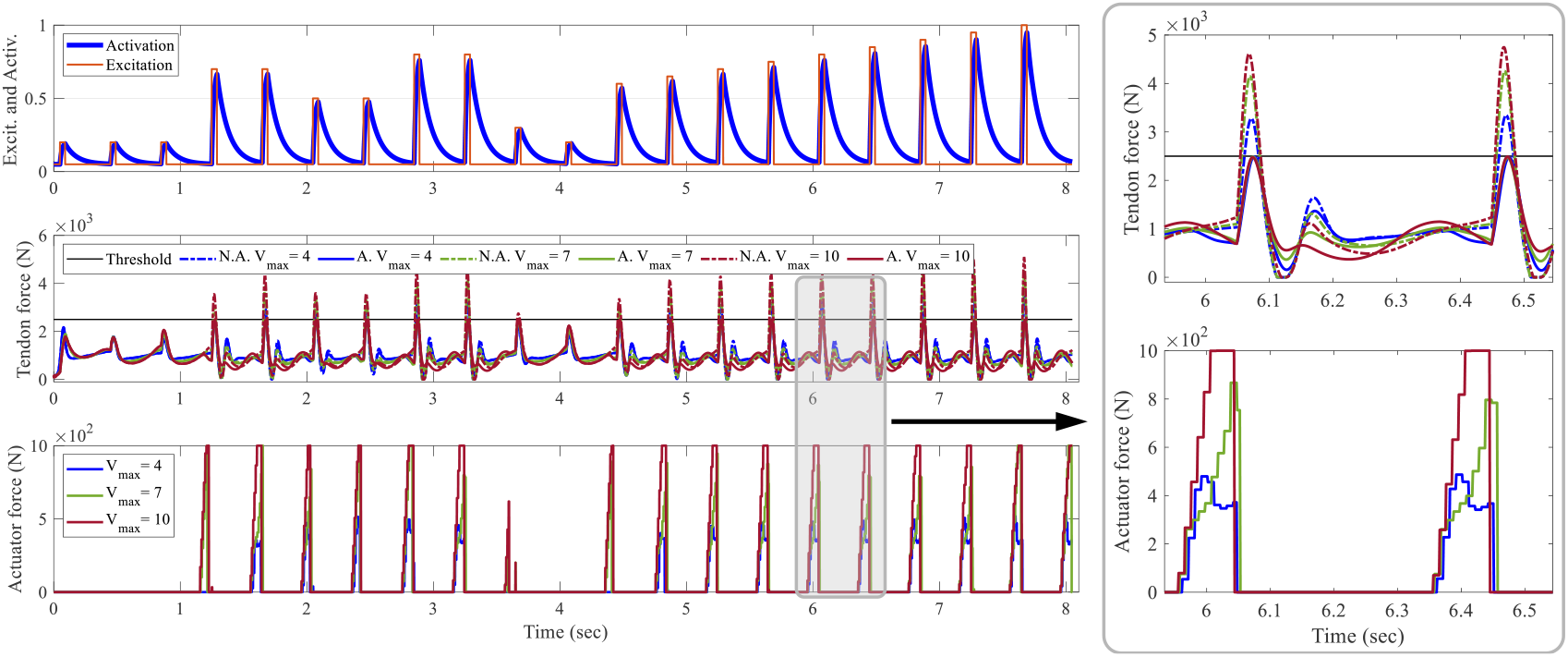
Predictive control performance applied to different muscle fiber types when the controller is aware of the fiber type.

#### 3.2.2 Robustness to changes in muscle fiber phenotype

For the three distinct muscle fiber phenotypes examined in this study, the controller effectively maintains the peak Achilles tendon force below the threshold. For the type I muscle fiber, the average peak tendon force, STD, and RMSE are 2471 *N*, 29.2 *N*, and 1.6%, respectively. These values are almost the same for the type II-A (and type II-X) as 2477 *N* (2482 *N*), 11.5 *N* (10.9 *N*), and 1% (0.8%). Furthermore, as observed, the non-assisted peak tendon force rises with the increase in the muscle fiber twitch velocity. Consequently, the faster twitch muscle fibers necessitate greater exoskeleton actuation to ensure that the assisted tendon force remains below the threshold.

The investigation into the robustness of the control framework was conducted without informing the controller of the changes occurring in the muscle fiber phenotype, see 6. In this scenario, the *V*^*max*^ parameters in both the main model and the controller were varied with different values. It was observed that when the *V*^*max*^ parameter in the controller exceeded its counterpart in the main model, the controller demonstrated some robustness by generating outputs to maintain the peak tendon force within the specified limit.

For instance, as illustrated in Figure 6, in the case where 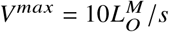 in the model and 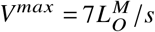 in the controller (represented by the dark red lines in Figure 6), the average peak controlled tendon force was 2783*N*. In this case, the RMSE and STD were calculated to be 12.2% and 116.5*N*, respectively. case where 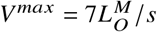 in the model and 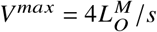 in the controller (represented by green lines in Figure 6), These statistics take the value of 2856 *N*, 15.5%, and 154 *N*, respectively.

**Figure 6.**
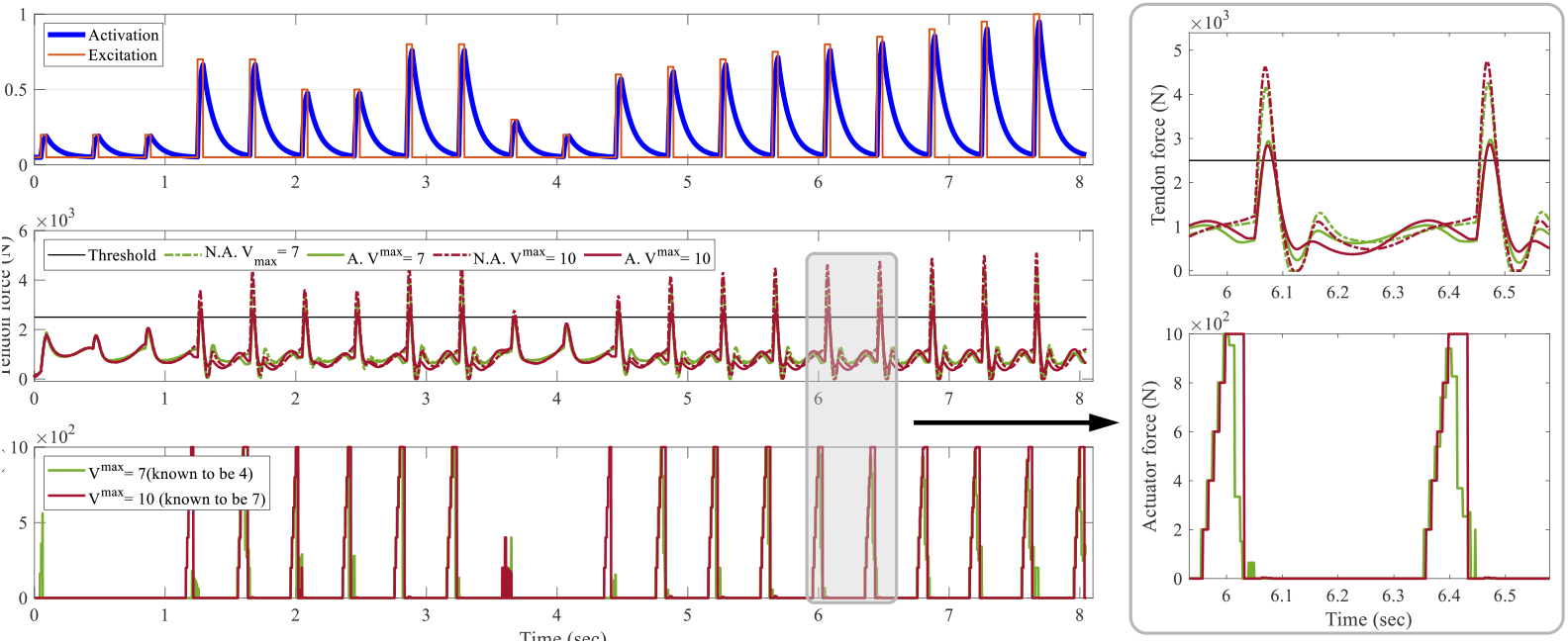
Robustness of NMPC to changes in muscle fiber type. The controller is not aware of the fiber type.

## 4. Discussions

In this paper, we introduced a closed-loop control framework designed to predictively control the peak Achilles tendon force during hopping via a parallel actuator. The proposed controller demonstrated robustness in delivering optimal assistive torques to the target MTU across a spectrum of assistance levels required by the user. Moreover, it exhibited the ability to adapt to the contractile speed charachteristics of different muscle fiber phenotypes ranging from slow to fast-twitch, showcasing resilience to variations in muscle type.

### 4.1. Modeling

When damping was incorporated into the conventional Hill-type muscle model, it became feasible to obtain a close-form solution for the tendon force in the form of ODE. As a result, the very low computational time of each frame of the model, 3*μs*, allowed us to use the model for predicting the future tendon force, knowing the muscle actuation of the future.

In the combined MTU-exoskeleton model, we made the assumption of a single lumped MTU instead of considering individual plantarflexing MTUs **[Ref]**. Additionally, we assumed a constant moment arm and that the exoskeleton actuation is applied at the same location as the Achilles tendon force. Future work will focus on extending and refining this model to incorporate more detailed representations of the MTU and varying moment arm dynamics.

### 4.2. Control

#### 4.2.1. Different prediction horizons and levels of robotic support

It was demonstrated that while small initial horizon values (*≤* 50*ms*) in the NMPC algorithm did not effectively constrain the peak Achilles tendon force below the threshold, they did result in a decrease from 5093*N* to 3671 *N*. As the initial horizon increased, the controller began activating earlier, mitigating the risk of surpassing the threshold and allowing for a more gradual increase in actuator activation. However, larger initial horizons, although more likely to maintain the peak tendon force within the limits, can lead to greater deviation from real-world applications in the presence of modeling uncertainties. Furthermore, larger control horizons result in increased computational time required for the controller. In all future simulations, the 100 *ms* initial horizon was implemented, as for the case of 75 *ms* the peak tendon force surpassed the threshold once (2505 *N*) and this might happen more often in hopping cases with higher frequencies. When possessing the 100 *ms* initial horizon, the computational time (14 *ms*) stays within the limits of MTU’s physiological electromechanical delay.

Gradually adjusting the support level offered by the exoskeletons can provide benefits. It can ais potential patients in regaining strength and mobility at their preferred pace, enhances the ability of elderly users to carry out daily activities with increased ease and independence, and assists individuals or athletes in optimizing their performance while mitigating the risk of fatigue-related injuries. Consequently, a crucial characteristic of the controller would be its robustness in delivering diverse levels of support without necessitating any modifications to the controller itself. By demonstrating an RMSE of 35.7 *N* for the peak MTU force and the threshold, the NMPC framework exhibited an acceptable performance in this aspect, highlighting its ability to provide adaptive assistance promptly.

#### 4.2.2. Changes in muscle fiber phenotypes

By accommodating diverse muscle phenotypes, we can extend assistance to a wider range of users with distinct physiological characteristics. The control of different muscle phenotypes enables the exoskeleton to enhance performance across various activities, spanning from rapid bursts of activity to sustained exertion. Additionally, due to the dynamic nature of muscle adaptation, transitional isoforms emerge as muscles possess the capacity to transition between these primary isoforms Wisdom et al. (2015). Consequently, having a controller that robustly adapts to changes in muscle phenotype over time would be advantageous.

While the controller was informed of the changes in the fiber types, it demonstrated robustness against various muscle fiber phenotypes by effectively maintaining the peak tendon force below and within a 1.2% margin relative to the threshold. However, when the controller operated without knowledge of the changes in the muscle fiber phenotypes and utilized muscle fiber types with slower twitch responses in the controller compared to the main model itself, it attempted to mitigate the amount of tendon force surpassing by actuating the exoskeleton. This was not the case when a muscle fiber with faster twich response was used in the controller.

As a limitation of our work, it should be noted that different muscle fiber phenotypes display variations not only in *V* ^*max*^ but also in 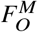. However, for the purposes of this study, our focus was specifically on adjusting the *V* ^*max*^ parameter while disregarding alterations related to 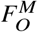.

In future work, the existing control framework will be deployed on a bowden-cable driven exoskeleton for both hopping and walking scenarios. In this setup, instead of relying on feedback solely from the main model (hopping model in the context of this study), the lengths of the musculotendon units (*L*^*MT*^) and tendon forces will be obtained in real-time from EMG and kinematic data collected from the user. These data will then be processed using the CEINMS real-time toolbox to derive MTU lengths and tendon forces.

## 5. Conclusion

In conclusion, this paper presents a pioneering framework for closed-loop control of peak tendon force within a simulated human ankle joint system equipped with parallel exoskeleton actuation. While previous research in lower limb wearable exoskeleton and exosuit control has primarily focused on reducing metabolic costs or compensating for biological joint torques, our work addresses the critical need for direct control over MTU dynamics. By integrating NMPC with a computationally efficient model encompassing Hill-type MTU contraction dynamics and ankle joint motion with parallel exoskeleton actuation, we bridge a significant gap in wearable robotic control.

Our approach demonstrates robustness across different levels of support and diverse MTU fiber phenotypes, from slow type I to fast type IIx, essential for functional movements like locomotion. By effectively controlling peak Achilles tendon force in various simulated conditions, including cyclic force productions in ankle plantarflexors, our framework lays the groundwork for future wearable robots capable of providing precise support tailored to specific MTUs. The computational efficiency of our model, with microsecond-level computation times and robustness to different muscle contraction velocities, underscores its potential for real-world applications.

## Supplementary material

## Ethical standards

### Author contributions

M.N, G.S.S, and M.S. conceived and designed the study. M.N. conducted all the simulations and performed the analyses and wrote the article. M.N. and M.S. reviewed and edited the article. M.S. ensured financial support for the work.

